# Understanding functional brain reorganisation for naturalistic piano playing in novice pianists

**DOI:** 10.1101/2024.01.15.575623

**Authors:** Alicja M. Olszewska, Maciej Gaca, Dawid Droździel, Agnieszka Widlarz, Aleksandra M. Herman, Artur Marchewka

## Abstract

Learning to play the piano is a unique complex task, integrating multiple sensory modalities and higher-order cognitive functions. Longitudinal neuroimaging studies on adult novice musicians show training-related functional changes in music perception tasks. The reorganisation of brain activity while actually playing an instrument was studied only on a very short time-frame of a single fMRI session, and longer interventions have not yet been performed. Thus, our aim was to investigate the dynamic complexity of functional brain reorganisation while playing the piano within the first half year of musical training.

We scanned twenty-four novice keyboard learners (female, 18-23yo) using fMRI while they played increasingly complex musical pieces after 1, 6, 13 and 26 weeks of training.

Playing music evoked responses bilaterally in the auditory, inferior frontal and supplementary motor areas, and the left sensorimotor cortex. The effect of training over time, however, invoked widespread changes encompassing the right sensorimotor cortex, cerebellum, superior parietal cortex, anterior insula and hippocampus, among others. As the training progressed, the activation of these regions decreased while playing music. Post-hoc analysis revealed region-specific time-courses for independent auditory and motor regions of interest. These results suggest that while the primary sensory, motor and frontal regions are associated with playing music, the training decreases the involvement of higher-order cognitive control and integrative regions, and basal ganglia. Moreover, training might affect distinct brain regions in different ways, providing evidence in favour of the dynamic nature of brain plasticity.

**Significance statement:** Mastering the piano is a unique process requiring a collaboration of multiple brain regions. However, associated brain activation changes are not well understood. Using functional MRI, we showed that playing the piano activated brain areas associated with movement and auditory processing. However, as the novice pianists progressed through their training, the activation of other brain regions, involved in memory retrieval, auditory-motor integration, or the processing of musical syntax, was gradually reduced. Our findings suggest that musical training is an optimisation process, with higher-order cognitive networks being activated more strongly at the beginning and their activity decreasing with increased proficiency.

## Introduction

Musical training has been frequently employed as a model to study neuroplasticity. Playing a musical instrument involves feedback and feed-forward mechanisms across multiple sensory and motor networks (Zatorre, Chen, and Penhune 2007; Brown, Zatorre, and Penhune 2015). Multiple cross-sectional studies demonstrated differences in brain structure and function between musicians and non-musicians in many brain areas, including the auditory cortex, the primary motor and premotor areas, parietal areas, and the white matter tracts connecting them (for reviews, see Zatorre, Fields, and Johansen-Berg 2012; Pantev et al. 2015; Bermudez and Zatorre 2005; Herholz and Zatorre 2012; Olszewska et al. 2021; Criscuolo et al. 2022). However, using cross-sectional studies alone, it is not possible to establish whether there is a causal link between musical training and observed differences between the two populations.

Longitudinal studies in adult learners show that musical training induces functional neuroplastic changes on timescales from minutes to weeks to months (for review, see Olszewska et al. 2021). Specifically, two single-session studies investigated playing the piano during a single fMRI session (Chen, Rae, and Watkins 2012; Brown and Penhune 2018). In those studies, a decrease in the activation of parietal, premotor and auditory regions was observed in late compared to early training, while the activation in the occipital cortex increased. Both studies have shown decreased activation of the auditory cortex for music perception tasks after training. However, these experiments were performed only in the initial (acute) phase of learning. Studies which considered longer training paradigms (lasting between 4 and 26 weeks) focused on adaptations in the resting-state networks or near-transfer effects related to music listening, not playing the instrument (Wollman et al. 2018; Herholz et al. 2016; Qiongling Li et al. 2018; Q. Li et al. 2019). Music listening is a relevant task in the context of musical training but does not provide insight into playing the instrument itself. In the current study, we focus on the changes in brain activity related to playing the piano in young adults undertaking piano training for the first time. This way, we shed light on the brain adaptations necessary to acquire a new skill.

Altogether, previous studies show evidence of functional training-related plasticity in the dorsal auditory stream, which is proposed to integrate sound and movement in musical tasks (for a recent review, see Olszewska et al. 2021, and a recent meta-analysis, see Pando-Naude et al. 2021). However, it is not clear how the brain activation while playing the piano changes when longer training is employed. First, it is proposed that the neuroplastic changes might follow an expansion-renormalisation pattern (Wenger et al. 2017), which cannot be captured on a short timescale or in a pre-post design. Indeed, studies on motor sequence learning point to differential patterns of brain activation between the acute and consolidation phases of training (see Doyon et al. 2009, and Penhune and Steele 2012, for reviews). Thus, we can expect that the time-course of changes in brain activation related to training might be region-specific and not uniform across the whole brain. Similar approach has been applied to understand the dynamics of training-related plasticity in linguisting processing (Matuszewski et al. 2020; Banaszkiewicz et al. 2020; Kuper et al. 2021). The musical training in the previous research was short compared to real-life conditions, where musicians practise for many years, and thus could not delve into the dynamic nature of temporal changes. Additionally, most fMRI studies use few, short musical pieces composed specifically for their purpose. Such experimental design choices certainly increase controllability of the experimental conditions, at the cost of real-world validity.

Thus, we investigated the time-course of brain reorganisation in novice pianists for the task of playing music over six months of naturalistic training. We specifically focused on the playing music task as opposed to near-transfer to other cognitive domains related to the auditory processing, which was the topic of many previous studies. We posed three research questions: (1) Which brain regions are involved in playing music in a naturalistic paradigm? (2) Can we observe dynamic changes related to musical training in brain activity during a naturalistic playing paradigm, and are the changes localised in the regions involved in playing music? (3) Are these changes uniform across the sensorimotor and auditory brain systems engaged in the playing task, or do they differ between regions of those brain systems?

We hypothesised that (1) the brain activation in the playback task should include the canonical auditory and motor regions; (2) training-related changes will encompass the regions of the motor network (motor cortex, pre/supplementary motor areas, striatum, cerebellum) and the dorsal auditory stream; (3) there would be a region-by-time interaction during training within three independent regions of interests (ROI) in auditory and motor areas associated with music production (Pando-Naude et al. 2021).

## Materials and Methods

### Participants

Previous longitudinal studies on piano-training-related neuroplasticity using fMRI had a sample size between 16 (Herholz et al. 2016) and 29 (Qiongling Li et al. 2018). Thus, we recruited a group of 24 healthy, right-handed female students (age range 18-23, M=21.3, SD=1.4) to undergo a 26-week piano training course. A recent methodological study estimated that sample sizes of around 25 participants are enough to represent average human functional brain organisation among groups (Marek et al. 2022). The participants were interviewed for involvement in musical practice, musical abilities, family history and education at the recruitment phase and to ensure that they were musically naïve, except for obligatory music classes in the Polish general curriculum for primary education. All participants were native Polish speakers, and had a normal or corrected-to-normal vision, normal BMI, lack of psychiatric and neurological illness, and unimpaired hearing. All participants provided written informed consent and were compensated for their time. This study was approved by the Research Ethics Committee at the Institute of Psychology of the Jagiellonian University, Kraków, Poland and the experiment has been carried out in accordance with The Code of Ethics of the World Medical Association (Declaration of Helsinki).

### Piano training course

The 26-week piano training course was conducted by a professional piano teacher using a unified, original approach. The curriculum, designed by Mrs Katarzyna Kiwior, encompassed learning 8 piano pieces of increasing complexity (Table 1), as well as technique basics, such as *staccato/legato*, technical motor exercises, and some gross and fine motor skill exercises. Participants were required to attend 13 (mostly) bi-weekly classes of 45 minutes, in pairs, either in person or remotely, depending on the current COVID-19 restrictions and health status. Deviations in the biweekly schedule occurred occasionally due to events such as serious illness and national holidays, such as winter and spring break, which were not included in the course duration. Each participant received their own electronic keyboard instrument for practising at home. A training schedule of around 30 minutes a day, for a total of about 4 hours a week, was advised. Moreover, the participants were required to attend a short, 15-minute video call with one of the experimenters (AO) every other week, when no piano classes were scheduled, to check on their progress and assert adherence to the training plan and time spent practising. An overview of the training course schedule can be found on the OSF (https://osf.io/q8abr).

**Table 1.** Piano training course material and playback task stimuli

| No | composer/source | title<br>(original title) | difficult<br>y | introduced | Measured<br>experimentally |
| --- | --- | --- | --- | --- | --- |
| M01 | Polish nursery rhyme | *A cat climbed a fence*<br>(*Wlazł kotek na płotek*) | easy | Week 1 | TP1, TP2, TP3, TP4 |
| M02 | French nursery rhyme | *Frère Jacques*<br>(*Panie Janie*) | easy | Week 2 | TP2, TP3, TP4 |
| M03 | Highlander folk song | *In a brick cellar*<br>(*W murowanej piwnicy*) | easy | Week 4 | TP2, TP3, TP4 |
| M04 | Polish nursery rhyme | *A train comes from afar*<br>(*Jedzie pociąg z daleka*) | easy | Week 6-7 | TP3, TP4 |
| M05 | American holiday song | *Jingle bells* | difficult | Week 8-9 | TP3, TP4 |
| M06 | Y. Tiersen | *Amélie - Nursery Rhyme from Another Summer*<br>(*Amélie - Comptine d'un autre été*) | difficult | Week 11-15 | TP4 |
| M07 | Kuyavian folk song | *Red Apple*<br>(*Czerwone Jabłuszko*) | difficult | Week 18-21 | TP4 |
| M08 | Ch. Petzold (formerly attributed to J.S. Bach) | *Minuet in G Minor from the Notebooks for Anna Magdalena Bach* | difficult | Week 18-21 | TP4 |

Reading score (sheet music) was out of the scope of the curriculum and the participants were asked to refrain from learning it on their own until the completion of the study, to avoid any interaction with language-related processing in the brain. The use of foot-operated pedals was not included in the course.

### Experimental Design and Statistical Analysis

The participants underwent 7 scanning sessions, during 4 of which (after 1, 6, 13 and 26 weeks of training - TP1, TP2, TP3 and TP4, respectively) they were asked to play in the MRI scanner, among other tasks. The playback task and the experimental conditions have already been validated in a group of trained musicians (Olszewska et al. 2023). In summary, playing was performed in a supine position in an MRI scanner, with an MRI-compatible MIDI keyboard (Olszewska et al. 2023) [Fig. 1] placed on a dedicated support stand, and elbows supported in a comfortable position with beads-filled cushions. The participants could not see their hands during the scan and were asked to refrain from trying to look at their hands or the keyboard, limit unnecessary movements, and concentrate on the visual cues.

**Fig. 1.**
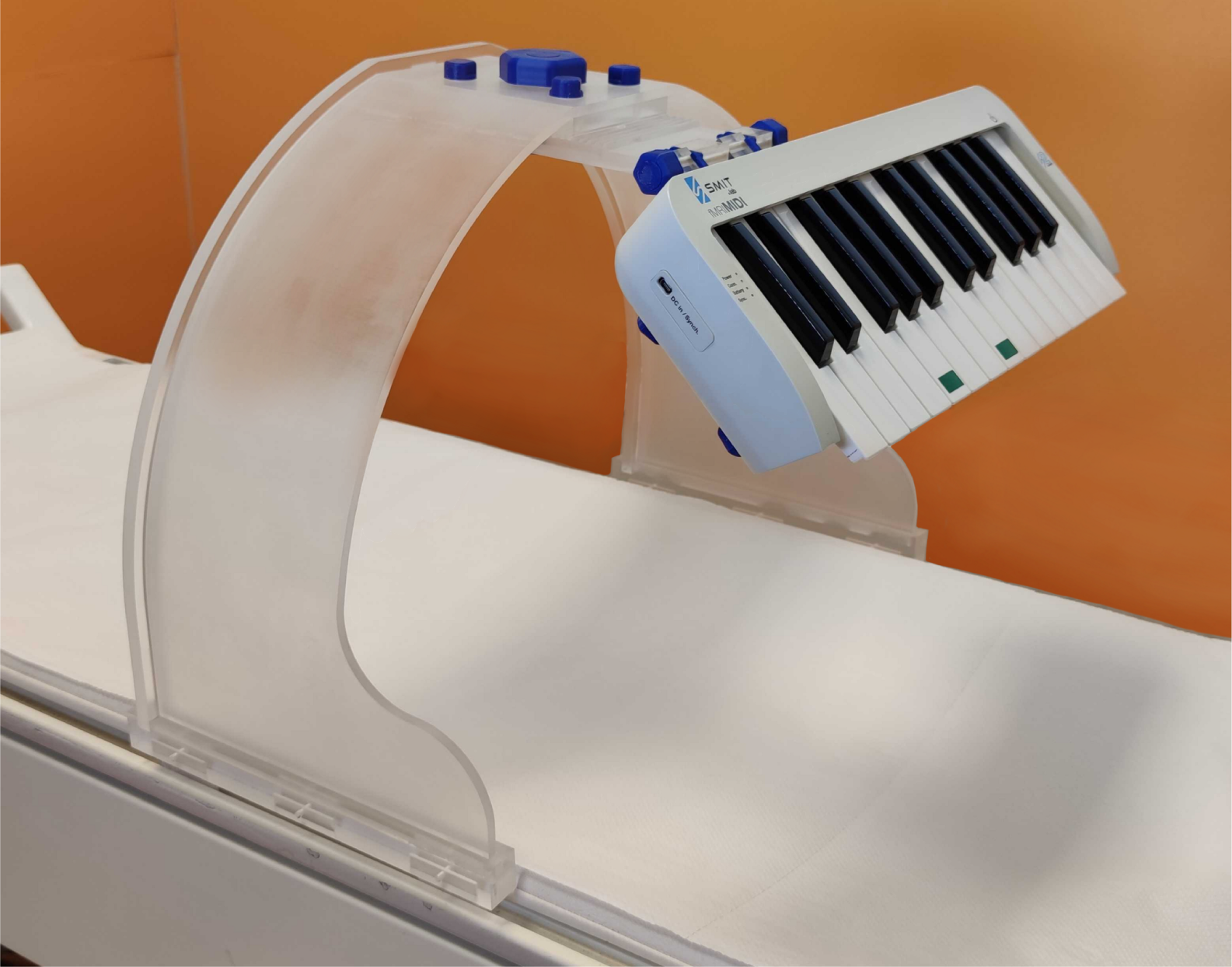
The MRI-compatible MIDI keyboard placed on a keyboard stand. Green convex stickers mark keys G3 and C4.

### Behavioural data

Behavioural data acquired during scanning encompassed the timing of key presses and releases for every key of the instrument. To investigate whether the participant’s learning resulted in improved performance, we scored the performances using a comparison to errorless performance by Levenshtein ratio and the relative rhythmic deviance, based on the tapping-PROMS paradigm (Georgi et al. 2022). The Levenshtein ratio is a method to compare sequences (Olszewska et al. 2023, Levenshtein and Others 1966) which relates the number of differences between sequences to their length. The Levenshtein ratio is defined as:

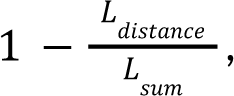

where Ldistance is the number of insertions, omissions and substitutions in the sequence performed by the participant compared to errorless performance, and Lsum is the sum of the lengths of the of both the performed sequence and the errorless performance (here, the sequence based on the score notation of the stimulus learned by the participant). Therefore, the Levenshtein ratio can be understood as the relative measure of how often the participant made a mistake while playing the target stimulus by measuring the similarity of the played sequence to errorless performance and yields a score between 0 (no similarity) and 1 (perfect accordance).

The relative rhythmic deviance was estimated using an approach based on tapping-PROMS, a method to compare rhythmic accuracy (Georgi, Gingras, and Zentner 2022, Olszewska et al. 2023). First, the relative duration of each note in the errorless performance is calculated with respect to its complete duration. For example, if the sequence expects four quarter-notes and two half-notes, the quarter-notes should last 12.5% of the whole sequence, and each half-note 25% of the total sequence duration. A similar transformation is calculated for the actual participants’ performance. In an actual performance, the notes might last 11.7%, 12.3%, 13.1%, 12.9%, 26.1% and 23.9% instead. The average relative rhythmic deviance is calculated:

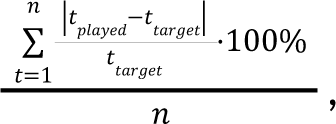

where tplayed are the relative durations of each the played notes, and ttarget their nominal durations, and n is the number of notes in the sequence. In this example, this yields 6.4%, 1.6%, 4.8%, 3.2%, 4.4%, 4.4% for each note, with a mean score of 4.3% deviation from nominal note duration. This score has a minimum of 0% and no maximal possible deviance score.

Statistical analysis of the behavioural data was performed using a generalised linear model in R (R Core Team 2020), including the effect of timepoint, the interaction between melody and hands condition, and the interaction between timepoint and hands condition. Subsequent visualisation was performed using Seaborn 12.0 (Waskom 2021) and matplotlib 3.7 (Caswell et al. 2023) in Python 3.10 (Van Rossum and Drake 2009).

### MRI data acquisition

Neuroimaging data were acquired on a 3-Tesla Siemens Magnetom Trio scanner with a 32-receive channel head coil. Functional data were acquired using echo-planar imaging pulse sequence in an Interleaved Silent Steady-State (ISSS) paradigm (Schwarzbauer et al. 2006), where 5 TRs (7.75s) of image acquisition were followed by 4 ‘silent’ TRs (6.2s) when the auditory stimuli were presented (multi-band acceleration factor 3, repetition time [TR]=1550 ms, echo time [TE] = 30.4 ms, flip angle [FA] = 56°). We acquired 60 slices in transverse plane orientation with an isotropic voxel size of 2.5 × 2.5 × 2.5 mm. For estimating magnetic field inhomogeneities, we additionally acquired two spin-echo EPIs with an inverted phase-encoding direction. An anatomical T1-weighted scan was acquired at the end of the scanning session using a magnetization-prepared rapid gradient-echo sequence (MPRAGE) with a voxel size of 1 × 1 × 1 mm isotropic (field of view = 256 × 176 × 256 mm [A-P; R-L; F-H]) in sagittal orientation.

### The Playback task

The task is similar to the playback task performed by professional pianists in a previous publication (Olszewska et al. 2023-playing music with naturalistic stimuli); however, the most difficult stimuli were replaced by simpler ones for the novices, yielding a set consisting of simple nursery rhymes and more difficult popular musical pieces, which were taught to the participants during the training course (Table 1). Briefly, at each timepoint, the participants would perform the material already mastered in the training course. At timepoint 1, this was a single simple melody, which was divided into two parts and played sixteen times, half with the right hand and half with both hands (*unisono*). To maintain task difficulty throughout the course, at each consecutive time point, the number and difficulty of melodies increased while the number of repetitions decreased, corresponding to participants’ progress in the course. The total task duration was kept at a constant of 32 trials occurring during the ‘silent’ 6.2s of the ISSS paradigm, each trial consisting of a listen and a playback phase interlocked with ‘scanning’ 7.75s of the ISSS paradigm. This resulted in each melody being played twice, once with the right hand and once with both hands, at TP4. The listen-playback design of the task was chosen to resemble how the participants learned their course material during classes, as no musical notation was introduced and the musical pieces were played by ear. Each melody fragment was presented to the participants aurally, and the participant was asked to recognise the melody and its fragment, and play it back after a short while. Auditory feedback was present during playing in the scanner task, however, participants could not see their hands, and their gaze was fixed on cues on a screen.

The overview of the stimuli used, the order and the number of repetitions at each timepoint can be found on the OSF (https://osf.io/q8abr).

### fMRI data analysis

At the subject level, fMRI data were preprocessed using a standard fMRIPrep pipeline, including fieldmap correction and excluding slice-time correction because of sparse-sampling acquisition [fMRIPrep 21.0.0 (Esteban, Markiewicz, Goncalves, et al. 2021; Esteban et al. 2019) RRID:SCR_016216, which is based on Nipype 1.6.1 (Gorgolewski et al. 2011; Esteban, Markiewicz, Burns, et al. 2021); RRID:SCR_002502].

The preprocessed functional files were then smoothed with a 6 mm FWHM Gaussian kernel within SPM12 (Wellcome Trust Centre for Neuroimaging, University College, London, UK, **Error! Hyperlink reference not valid.** running on MATLAB 2019b (MathWorks, http://www.mathworks.com). The analyses were performed using a general linear model (Friston et al. 1994) on a whole-brain level. To correct for sparse acquisition, the first acquired volume was used as a dummy scan to fill in the ‘silence’ gaps, a ‘0’-vector to expand the head movement regressors (translation and rotation in X, Y and Z directions), and an additional regressor was added in the model to indicate which scans were actual scans and which were the dummy scans (Peelle 2014).

On the first level, contrast images (playback vs. global baseline) were computed to provide task-related activity for each participant at each timepoint. Participants’ movement was analysed for excessive motion (framewise displacement > 7.5) and framewise displacement peaks were regressed out. Second-level group analyses were performed in a flexible factorial model including time and subject factors. To correct for multiple comparisons, a cluster-level height threshold of p < 0.05 (FWE), together with a cluster-level extent threshold of 5 voxels were applied. Contrast for the Main Effect of Task and Main Effect of Time were exported from SPM and visualised using nilearn (Abraham et al. 2014) in Python 3.10 (Van Rossum and Drake 2009). In order to verify the overlap between the main effects of time and task, a conjunction analysis was performed using the easythresh_conj script (Nichols 2007), which is a part of the FSL package (Smith et al. 2004). Anatomical localization was determined using the AAL3 atlas in bspmview (Spunt 2016) for peaks at least 8mm apart. For visualisation purposes and independent ROI analysis, time-courses from clusters’ peaks and ROIs were extracted using MarsBaR (Brett et al., n.d.) version 0.45 running on SPM12 and MATLAB 2019b.

### Independent regions of interest

To investigate the differences in time-courses between regions belonging to different brain networks, we used independent regions of interest including the right auditory and the left and right motor regions. Three music-production related regions from data published by (Pando-Naude et al. 2021): the *left sensorimotor* ROI, including the left postcentral and precentral gyri [MNI -43_-22_46]; the *right motor* ROI, including the precentral gyrus [MNI 53, 6 30]; and the *right auditory* ROI, including planum temporale, extending into the superior temporal gyrus and Heschl’s gyrus [MNI 59 -21 7] were downloaded from Open Science Framework. Afterwards, for each region of interest, we extracted first-level contrast estimates using MarsBaR (Brett et al., n.d.) version 0.45 running on SPM12 and MATLAB 2019b. The interaction between region and time was then analysed using two-way repeated-measures ANOVA in the Pingouin v0.5.1 package (Vallat 2018) in Python v. 3.10 (Van Rossum and Drake 2009). Subsequent visualisation was performed using Seaborn 12.0 (Waskom 2021) and matplotlib 3.7 (Caswell et al. 2023) in Python 3.10 (Van Rossum and Drake 2009). version 0.45 running on SPM12 and MATLAB 2019b. The interaction between region and time was then analysed using two-way repeated-measures ANOVA in the Pingouin v0.5.1 package (Vallat 2018) in Python v. 3.10 (Van Rossum and Drake 2009). Subsequent visualisation was performed using Seaborn 12.0 (Waskom 2021) and matplotlib 3.7 (Caswell et al. 2023) in Python 3.10 (Van Rossum and Drake 2009).

### Code Accessibility

The original code used for the analysis and visualisation of the data can be found on OSF https://osf.io/q8abr

## Results

### Behavioural performance

We analysed the behavioural performance of the participants to assure that the training was successful, reflected in improved performance over time. For melodic performance, a generalised linear model revealed a significant effect of time (t=2.360, p=0.018), a significant effect of hand condition (right hand/both hands) (t=6.022, p<0.001), and their interaction (t=-2.147 p= 0.032). Overall, the melodies performed with both hands were generally played at a lower performance, which improved over time, compared to melodies performed with only the right hand which had a stable performance in time. Moreover, more difficult melodies, which were introduced later in the course, were played at a lower performance compared to melodies introduced earlier in the course (Fig. 2). Similarly, for the rhythmic performance, we observed a significant effect of time (t=-2.419, p=0.015), hand condition (t=4.663, p<0.001), but not their interaction (t=1.763, p=0.078) (Fig.2.).

**Fig. 2.**
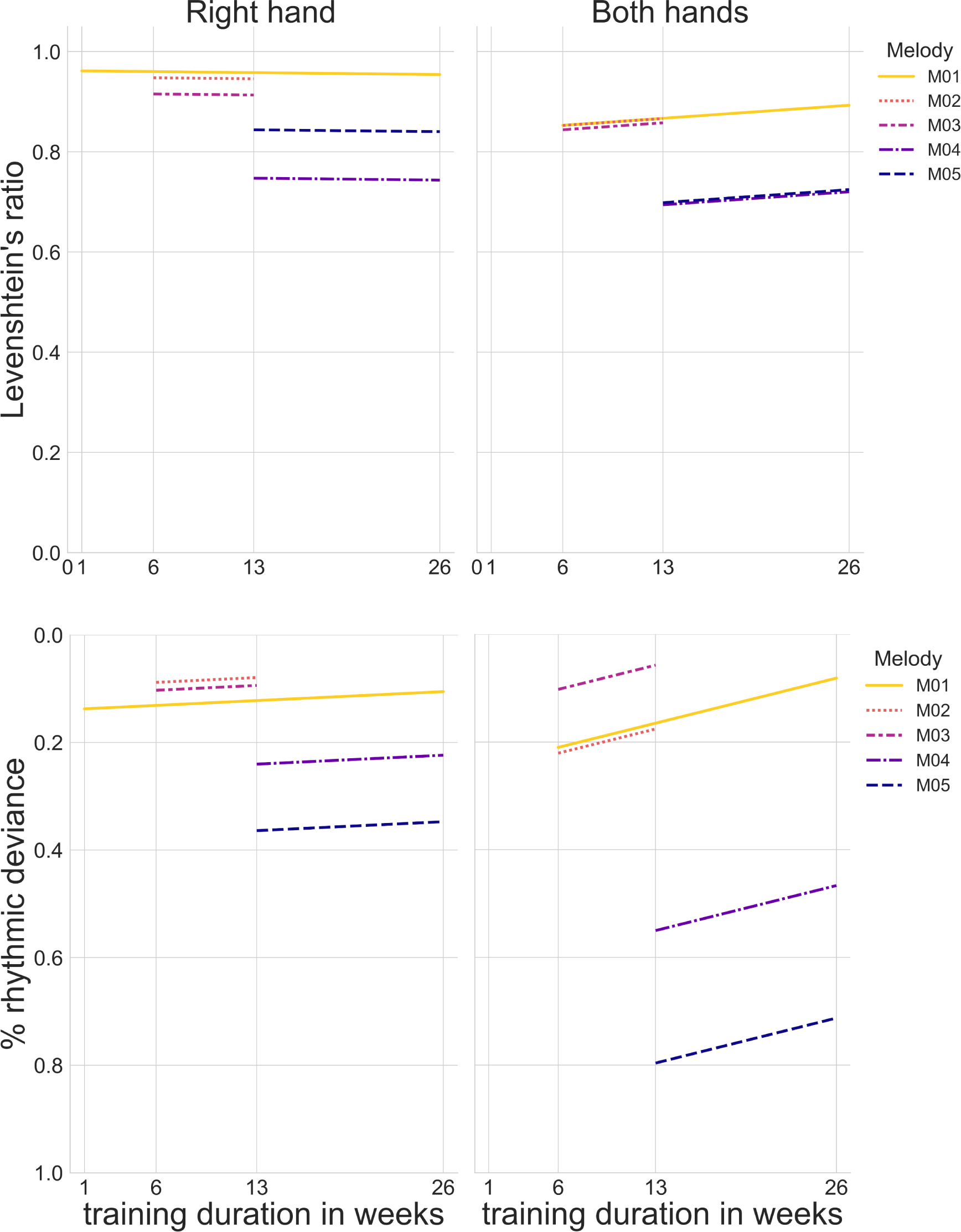
Behavioural performance for repeated melodies played with the right hand or with both hands. Top panel: melodic accuracy using Levenshtein ratio, bottom panel: rhythmic accuracy using percentage deviance from expected note duration. The data is presented in such a way that higher accuracy is always shown at the top of the plots.

### Neuroimaging results

In the playback task, we investigated the brain activity and the time course of brain reorganisation in novice pianists while playing the piano using naturalistic musical stimuli. One participant presented excessive motion at TP2 (6 weeks of training). The volumes corresponding to the excessive motion were regressed out.

### Main Effect of Task

Irrespective of time, the playback task revealed brain activation in the left precentral and postcentral gyri; the right temporal pole extending into the inferior frontal gyrus (pars opercularis) and the planum temporale; the IV, V and VI lobules of the vermis; the left superior temporal gyrus encompassing the left planum temporale; and bilaterally in the supplementary motor cortex. A summary of the brain activation related to the playback task can be found in Table 2 and Fig. 3A.

**Table 2.** Main Effect of Task. Local maxima separated by more than 8 mm. Regions were automatically labelled using the AAL3 atlas. X, Y, Z - Montreal Neurological Institute (MNI) coordinates in the left-right, anterior-posterior, and inferior-superior dimensions, respectively. R, L - right, left hemisphere.

| Main Effect of Task |  | MNI coordinates |  |  |  |
| --- | --- | --- | --- | --- | --- |
| Region Label (AAL3) | T-value | X | Y | Z | extent |
| L precentral gyrus | 13.41 | -27 | -25 | 74 | 243 |
| R temporal pole | 11.70 | 54 | 15 | -4 | 222 |
| Vermis IV V | 9.81 | 1 | -48 | -4 | 10 |
| L postcentral gyrus | 9.04 | -49 | -35 | 59 | 48 |
| L superior temporal gyrus | 8.52 | -59 | -30 | 14 | 126 |
| L supplementary motor area | 7.49 | -7 | 0 | 74 | 54 |
| Vermis VI | 5.84 | 1 | -75 | -11 | 5 |

**Fig. 3.**
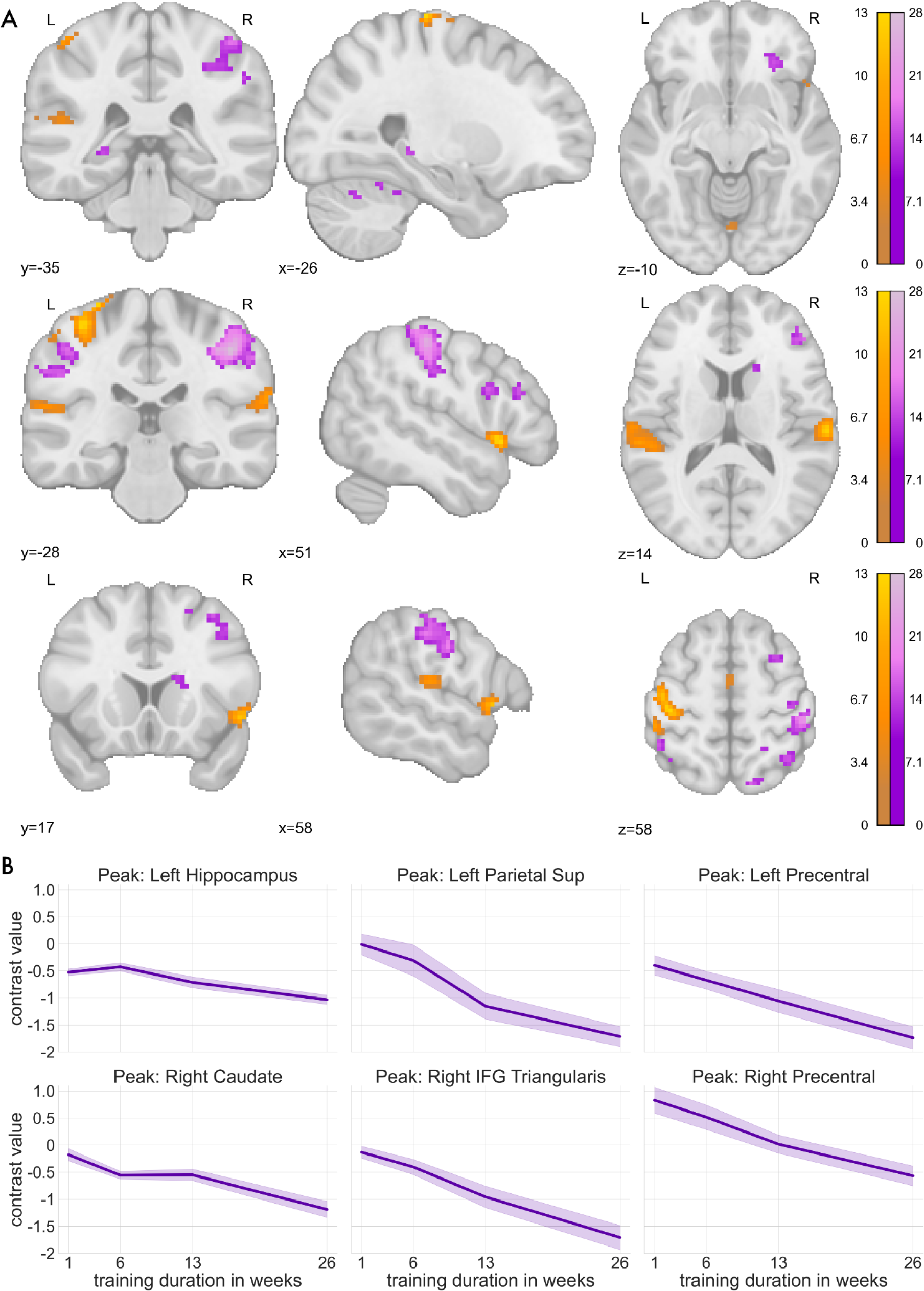
(A) Statistical maps representing the contrasts for the Main Effect of Task (orange) and Main Effect of Time (purple). x, y, z - MNI coordinates. Colour bars represent test-statistic ranges. (B) Visualisation of musical-training-related changes in brain activation in selected cluster peaks.

### Main Effect of Time

Musical training-related brain activation changes for the playback task encompassed multiple brain regions (Table 3, Fig. 3A), such as the right postcentral gyrus, extending into the precentral and supramarginal gyri, and the parietal cortex; the right inferior frontal gyrus, extending into the frontal pole and the middle frontal gyrus; the right caudate nucleus; bilaterally in the crus I, VI, VI and VIII of the cerebellum; the left precuneus; the left inferior temporal gyrus (at the temporo-occipital junction); the left parietal cortex; or the left hippocampus, among others. For visualisation purposes, we present the time-courses of selected cluster peaks (Fig. 3B). Pairwise comparisons between timepoints showed a significant effect for TP1 (1 week) > TP4 (26 weeks), TP1 (1 week) > TP3 (13 weeks) and TP2 (6 weeks) > TP4 (26 weeks) (Fig.4).

**Table 3.**
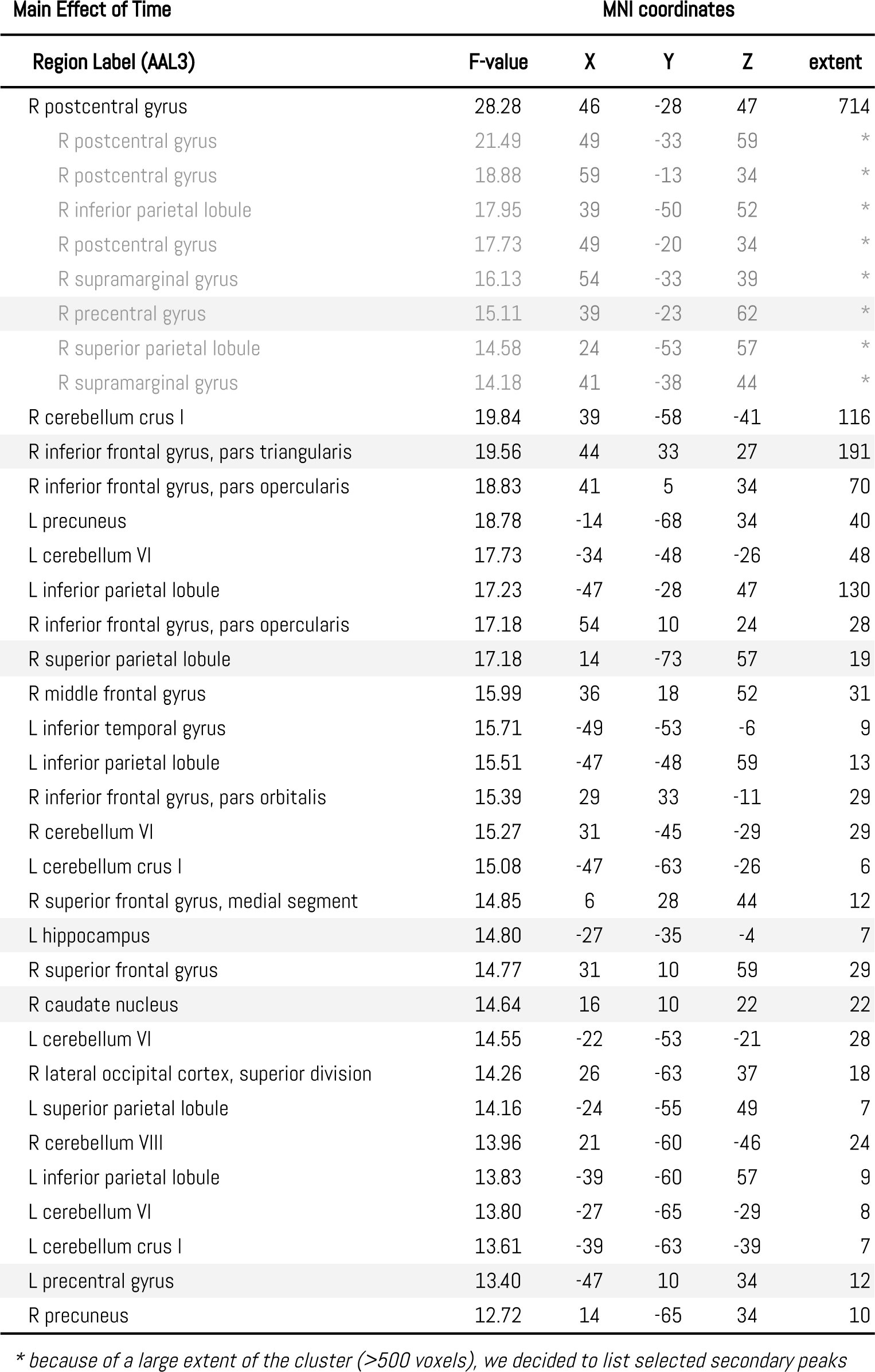
Main Effect of Time. Local maxima separated by more than 8 mm. Regions were automatically labelled using the AAL3 atlas. X, Y, Z - Montreal Neurological Institute (MNI) coordinates in the left-right, anterior-posterior, and inferior-superior dimensions, respectively. R, L - right, left hemisphere. Highlighted peaks were later used for the visualisations (Fig. 3B).

**Fig. 4.**
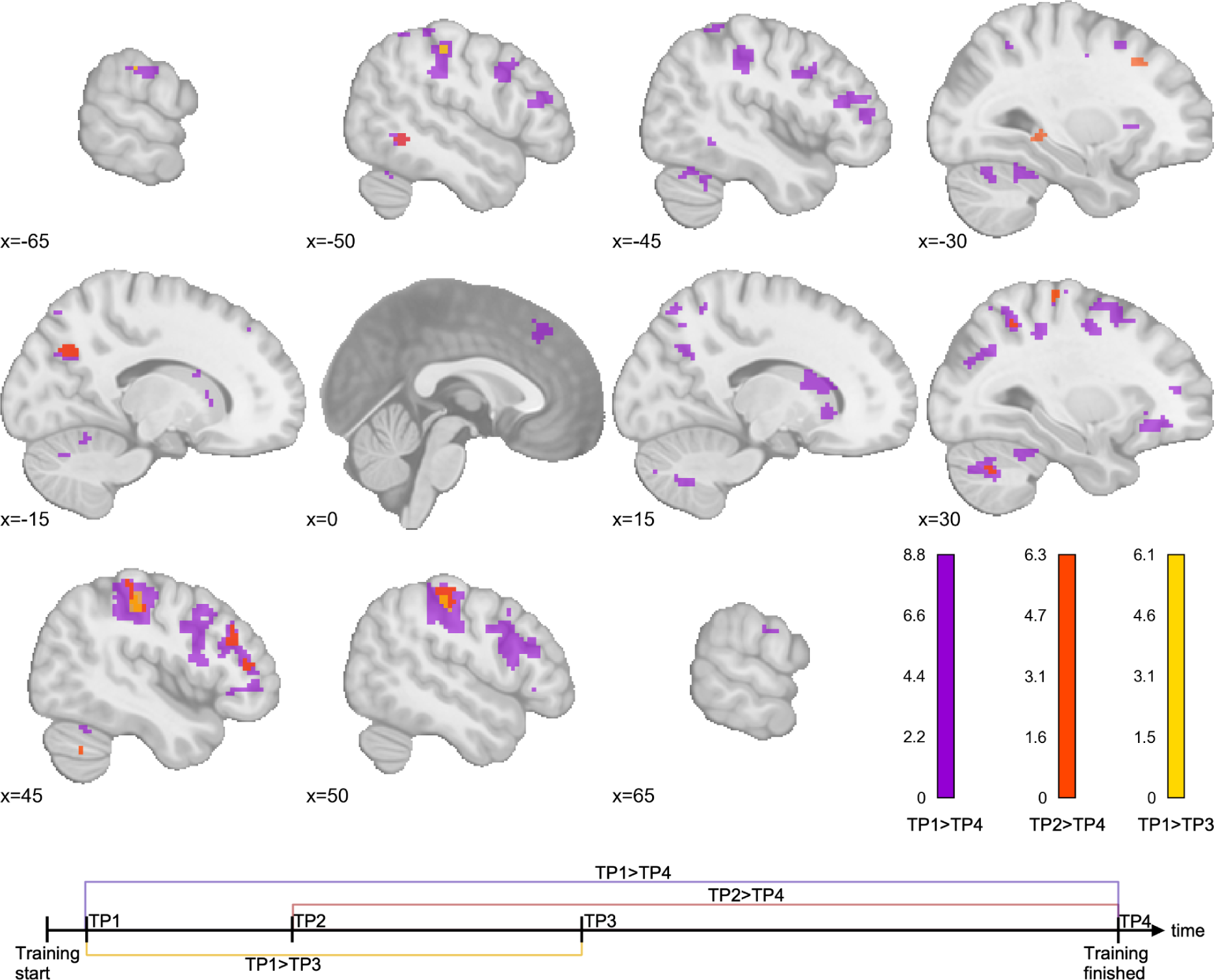
Pairwise comparisons between TP1 (1 week) > TP4 (26 weeks), TP2 (6 weeks) > TP4 (26 weeks) and TP1 (1 week) > TP3 (13 weeks). The remaining pairwise comparisons yielded no significant clusters. Colour bars represent test-statistic ranges.

### Independent Region of Interest (ROI) analysis

The independent ROI analysis between the *left sensorimotor* ROI, the *right motor* ROI and the *right auditory* ROI revealed significant effects of time, region and their interaction (Table 4, Fig. 5). In the pairwise comparisons analysis, there were significant differences between the following: TP1 (1 week) and TP4 (26 weeks) for the *left sensorimotor* ROI (t=4.521, p-corr=0.003) and *right motor* ROI (t=7.542, p-corr<0.001); TP2 (6 weeks) and TP4 (26 weeks) for the *right motor* ROI (t=4.378, p-corr=0.004); the *left sensorimotor ROI* and the *right auditory* ROI at TP3 (13 weeks) (t=-3.346, p-corr=-0.034); the *right auditory ROI* and the *right motor ROI* at TP3 (13 weeks) (t=4.550, p-corr=0.002); and, at the TP4 (26 weeks), all three ROIS (*left sensorimotor* ROI and *right auditory* ROI t=-5.048, p-corr<0.001; *left sensorimotor* ROI and *right motor* ROI t=4.815, p-corr<0.001, *right auditory* ROI and *right motor* ROI =7.263, p-corr<0.001).

**Table 4.** The results of repeated-measures ANOVA for the main effects of time, ROI and their interaction of training-related changes in independent ROIs. p-GG-corr: Greenhouse-Geisser corrected p-value; ηp2 - generalised eta-squared effect size.

| | df1 | df2 | F | p-GG-corr | effect size ($\eta^2$) |
| --- | --- | --- | --- | --- | --- |
| Timepoint | 3 | 69 | 5.225 | 0.004 | 0.185 |
| ROI | 2 | 46 | 13.924 | <0.001 | 0.377 |
| Timepoint * ROI | 6 | 138 | 15.483 | <0.001 | 0.402 |

**Fig. 5.**
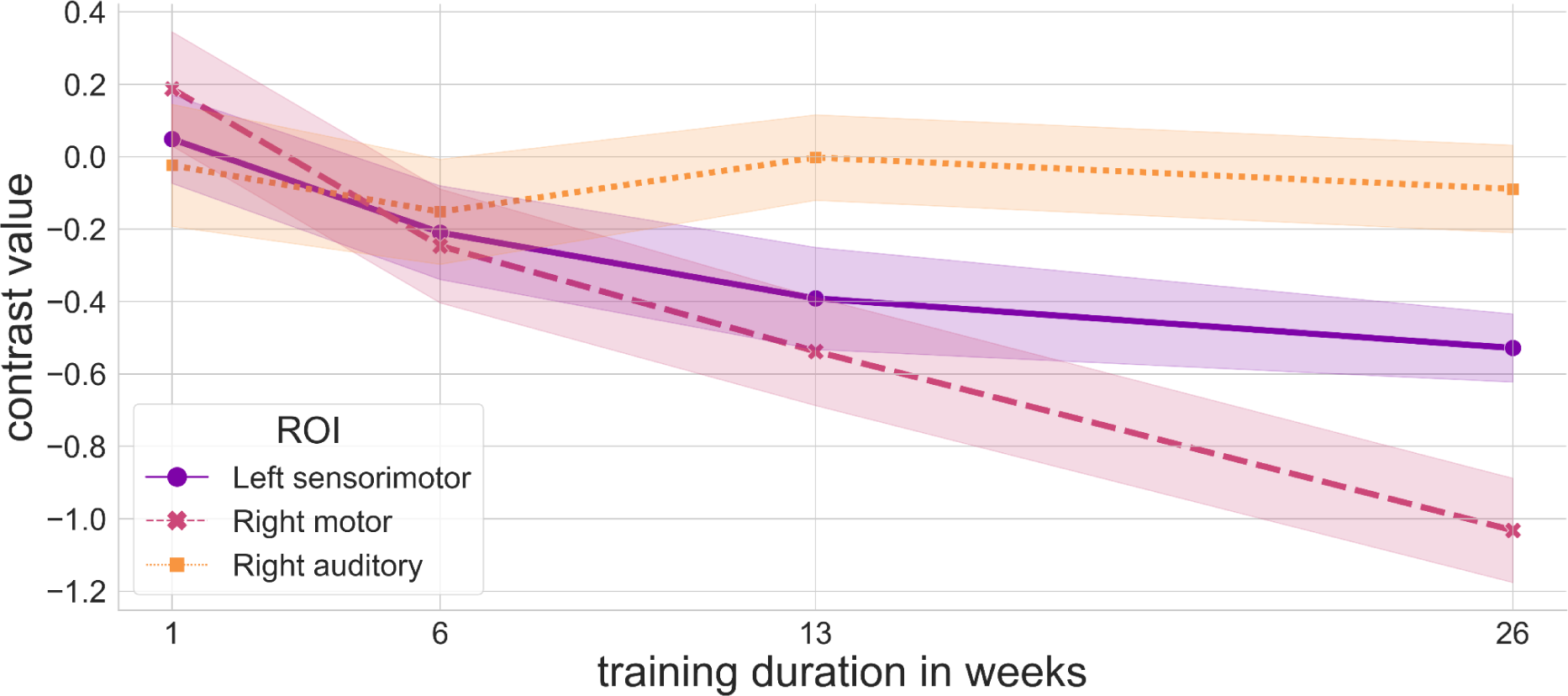
Visualisation of musical-training related changes in brain activation in independent ROIs.

### Conjunction Main Effect of Task ⋂ Main Effect of Time

The conjunction analysis of the main effects of task and time revealed no significant clusters, and therefore is not presented in the tables or visualisations.

## Discussion

In this study, we used fMRI methodology to investigate the dynamics of neuroplastic changes related to learning to play a keyboard instrument for 6 months. We used an originally developed naturalistic paradigm for both the training and the experimental tasks, with increasing difficulty as the training progressed. Our findings show that, over the course of training, the performance of stimuli played with both hands improves with time (Fig. 2.). The main neuroimaging findings include: (1) the main effect of task (playing music) activates the canonical auditory-motor network related to musical performance (Fig. 3A.); (2) the main effect of time reveals training-related changes in multiple regions related to motor control, memory or higher-order cognitive control (Fig. 3AB); (3) there is a region-by-time interaction of neuroplastic changes in selected independent regions of interest (Fig 5). We discuss the interpretation and significance of each of the findings below.

### The performance of stimuli played with both hands improves with time

We show that the performance of stimuli, measured as a similarity to errorless performance, improves both for the order of keys pressed, as well as for rhythmic accuracy (Fig.2). The stimuli played with the right hand only are generally played better than stimuli played with both hands, and the more complex stimuli introduced later in the training present overall lower performance compared to stimuli introduced at the beginning of the course. The lower production of the stimuli played with both hands and more difficult stimuli might be related to their complexity, or the overall number of stimuli present at each timepoint (starting from excerpts from a single melody at TP1 compared to playing excerpts from all 8 melodies at TP4). Nevertheless, we interpret that the increased performance of stimuli with time reflects the learning process and the improved ability to play the instrument.

### (1) The playback task reveals the canonical auditory and motor regions involved in playing music

Averaging across timepoints, we identified a canonical network related to playing a musical instrument, including the bilateral auditory and left motor, as well as supplementary motor and cerebellar regions (Fig.3A). This is in line with previous studies on music production (for reviews, see (Zatorre, Chen, and Penhune 2007; Pando-Naude et al. 2021). The supplementary motor area is one of the regions taking part in visuomotor coordination and learning (Kawashima et al. 1994). The occurrence of the left but not right sensorimotor regions might be caused by half of the trials being right-hand only, or the right-hand dominance of the participants. Apart from the auditory and motor regions, we also observed activation in the bilateral temporal poles and the inferior frontal gyrus (pars opercularis). The temporal poles are multifunctional regions involved in many cognitive functions, including naming and word-object labelling or semantic processing in all modalities (Herlin, Navarro, and Dupont 2021). We postulate that the involvement of temporal poles in playing music in the current task could be related to these aspects, as the task included recognition of the melodies prior to playing them. In previous studies, the difficulty in recognition and naming of musical pieces was associated with damage to temporal poles (Johnson et al. 2011; Belfi and Tranel 2014; Belfi, Kasdan, and Tranel 2019; Schneider et al. 2018). Alternatively, one study linked the activity in the temporal pole to decoding the temporal structure of music and speech (Abrams et al. 2011). To avoid concluding from reverse inference, further study could investigate the exact relationship between the activity in frontal poles and playing and recognition of known musical excerpts.

The pars opercularis is a part of the inferior frontal gyrus which, in the context of speech and music, is proposed to represent syntactic and phonological processing as well as motor control. Altogether, the task allowed us to properly identify the brain areas involved in playing a musical instrument in novice pianists and supports the feasibility of the task in novices.

### (2) Musical training induces changes in the engagement of regions involved in cognitive and motor control during piano playing

The Main Effect of Time revealed widespread training-related changes in multiple brain networks (Fig.3A). These changes include, among others, regions involved in procedural movements and motor planning (caudate nucleus, cerebellum), musical syntax processing and error monitoring (inferior frontal gyrus), memory retrieval (posterior hippocampus), and auditory-motor integration (superior parietal lobule). We observed a decrease in activation with time in all regions, as visualised by peak activation trajectories (Fig.3B) and revealed by pairwise comparisons between timepoints (Fig.4). Additionally, there was no overlap between the regions revealed in the Main Effect of Task and the Main Effect of Time. These results are consistent with research on simple motor training, where a decrease in the activation of parietal and premotor cortices was associated with training, in a disjunction from regions involved in the task itself (Garzón et al. 2023). A decrease in the activation of motor-related areas during motor training has been previously shown in human and animal model studies (for review, see Wenger et al. 2017), apart from the very initial period, which we were not able to include for practical reasons, as discussed in the limitations. A decrease in brain activation in the late compared to early training was also shown in the two single-session studies where novices learned to play simple melodies (Chen, Rae, and Watkins 2012; Brown and Penhune 2018). Yet, these results need to be interpreted carefully, as the decrease in activation within a single session might reflect a repetition-suppression effect (Garrido et al. 2009; Brown et al. 2013). In a study comparing drummers with long-term training and non-musicians on an audiovisual asynchrony detection task, it was observed that musical training expertise is associated with reduced brain activation in the cerebellum, parahippocampal gyrus, inferior parietal lobule, inferior and middle temporal gyri, and precentral gyrus when performing the task (Petrini et al, 2011). A decreased activation might reflect an underlying optimisation process, where training results in reduction of neural processing involved in task performance. However, another study (Lee and Noppeney 2011) revealed increased activation in pianists compared to non-musicians in the STS-premotor-cerebellar circuitry when performing a similar task.

In the current study, we see an asymmetry in training-related activation decrease of the sensorimotor areas related to the movement of the left but not the right hand on the whole-brain level. This might be related to the right-handedness of the participants. In right-handed individuals, precise movements of the fingers of the right hand are already established by early adulthood, while such movements of the left hand are more demanding (Gut et al. 2007; Lee, Jin, and An 2019). These results are supported by the outcome of the independent regions of interest analysis.

### (3) Region-dependent time courses are identified in three regions of interest involved in music production

Finally, we observed a region-by-time interaction between three regions of interest associated with music production, identified in the independent meta-analysis (Pando-Naude et al. 2021), including the *right auditory*, *right motor* and *left sensorimotor* regions. There is no change in time in the auditory ROI, while both motor-associated ROIs show a significant decrease in activation at the final timepoint compared to early training. After 13 weeks of training, there was a significant difference between the auditory and both of the motor ROIs. After 26 weeks, there was a significant difference between all three ROIs, including a difference between the left and right motor regions, with the activation in the right motor region, associated with left-hand movement, decreasing more steeply than the activation in the left motor region. This indicates that, during the first 6 months of music production training, auditory and motor regions undergo functional reorganisation with distinct dynamics. While the activity of auditory regions remains relatively unchanged, the *right motor* regions undergo changes at the fastest rate, and the *left sensorimotor* region changes at a slower pace. These results are in line with the interpretation based on the whole-brain analysis, that brain activation related to left-hand movement requires more fine-tuning than the right-hand in these participants.

Contrary to previous longitudinal studies (Che et al. 2022, Lappe 2008, Lappe 2011), we do not observe any adaptations in the auditory cortices of novice pianists as they gained proficiency. This also stands in contrast to studies showing differences in auditory processing between musicians and non-musicians (e.g. Schmithorst and Holland 2003, Angulo-Perkins et al. 2014). While the absence of evidence does not equal evidence of absence, this discrepancy suggests alternative explanations. First, it is tempting to point out that the half-year of training, while long for research standards, is not enough to reflect changes observed in cross-sectional studies comparing musicians and non-musicians. However, the previous longitudinal studies we mention used even shorter training paradigms, in the order of several weeks at most.

Alternatively, perhaps the adaptations occur very early in the training and stabilise later on. In the current study, we could only start performing measurements after the first week of training, therefore if the changes in auditory processing occur specifically within this first week, they could not be detected with current design.

Although recruiting a control group was not feasible due to task design in the current study, in a recent resting-state connectivity study (Herman et al 2023) we documented that the functional connectivity did not change between or within any of the brain networks in the training group during the whole study period. This may suggest that the differences in brain activity observed in previous task-based fMRI studies might reflect the salience of the stimuli and not a difference in their sensory processing. For auditory processing, salience and attention are strong modulators of sensory perception, which is reflected in the well-known cocktail party effect. As such, the differences in results might also stem from differences in the tasks themselves. In the current task, the primary focus was playing music while listening to auditory feedback was an implicit element of this task. If the salience of the stimuli and attention processes regulate the activation in the auditory cortex, plausibly the involvement of the auditory cortex is increased only for explicit auditory listening tasks but not implicit ones such as the playing music task described here. Therefore, we argue that increased brain activation in the auditory cortices while performing explicit music listening tasks might reflect changes related to attention, and not sensory processing itself. Task context has been demonstrated to modulate brain activation in an EEG music listening experiment (Markovic et al. 2017) and functional connectivity in an fMRI experiment on emotional and reward processing in music (Liu et al. 2017), and there is evidence of top-down regulation of early auditory MEG responses to ambiguous rhythm (Iversen et al. 2009).

### Limitations

The strength of the current study lies in its naturalistic design, validated methodology and multiple time points revealing the dynamic nature of neuroplastic changes. However, no research is without limitations, and there are certain aspects which need to be taken into account when interpreting the results of current study. First, since we were interested explicitly in the task of playing the piano, this task could not be performed by completely naïve individuals, such as participants before the start of the training or a control group. Therefore, we were not able to investigate any changes within the first week of training, or perform a comparison to a control group. Yet, in a previous paper, we have compared the resting-state functional connectivity patterns of the novice pianists training group with that of trained pianists (Herman et al, 2023). We showed that no measurable changes in interoceptive ability or resting-state connectivity occurred within the study timeframe. A second limitation is that the participants could not look at their hands while playing in the scanner. However, the participants were familiar with this limitation and practised playing without looking at their hands at home. Additionally, the age range of participants is relatively narrow, spanning what can be considered young adulthood. As such, current results cannot be easily generalised and compared to studies on children or elderly, as developmental and ageing processes might interact with the experience-based neuroplastic processes studied here.

Finally, various aspects of auditory cognition have been found to differ between musicians and non-musicians, including emotional processing of music (Martins, Pinheiro, and Lima 2021), speech perception (Neves et al. 2022), multisensory integration (Landry and Champoux 2017, Che et al. 2022), or action-perception coupling (Novembre and Keller 2014). These are not covered here but are important aspects to study in future longitudinal research on novices to establish causal links between music training and the differences observed in cross-sectional studies.

### Conclusions

In summary, the current study, featuring a distinctive longitudinal design encompassing multiple time points, offers fresh insights into the dynamics of experience-induced neuroplasticity. Using a naturalistic paradigm, we show that piano training in young adults causes a decrease in activation of motor-associated and integrative brain areas while playing the instrument, similarly as previously identified in very short piano trainings (Chen et al, 2012, Brown & Penhune, 2018) and studies on non-musical motor training (Garzon et al, 2023). Neuroplastic responses in the activation of brain structures engaged in piano training are region- and timeframe-specific. For example, the changes in the striatum and premotor cortex are observed from week one, whereas the hippocampus starts to respond only after six weeks of training. This is also illustrated in the ROI analysis, which shows differential time-courses for the left and right (sensori)motor regions. To our knowledge, this type of patterning of brain reorganisation in space and time, previously described for motor training (Doyon et al. 2009, Penhune and Steele 2012) has not been investigated in the context of musical training. By uniquely employing multiple timepoints and fMRI music playing task, we illustrate not only the extent, but also the time-course of neuroplastic changes.

We observe that longer training results in more widespread changes in the brain. However, this is not characteristic of the brain plasticity model of map expansion, as the brain activity in those regions decreases, and the areas identified to be primarily activated by the task remain unchanged. This pattern is indicative of the optimisation of higher-order brain areas involved in motor control and auditory-motor integration as well as memory and the processing of musical syntax for the task of playing the piano. Such reorganisation is in line with the latter phase in the ‘expansion-renormalisation’ model of neuroplasticity (Reed et al. 2011), where, after the initial period, training results in the stabilisation of efficient brain circuitry, which supports the learned behaviour.

## Author information

### Alicja M. Olszewska

- Laboratory of Brain Imaging, Nencki Institute of Experimental Biology of the Polish Academy of Sciences, Warsaw, Poland **Contribution:** Conceptualization, Data curation, Software, Formal analysis, Validation, Investigation, Visualisation, Methodology, Writing - original draft, Writing - review and editing, Project administration

**Competing interests:** No competing interests declared ORCID 0000-0002-3995-8166

### Maciej Gaca

- Laboratory of Brain Imaging, Nencki Institute of Experimental Biology of the Polish Academy of Sciences, Warsaw, Poland

**Contribution:** Data curation, Software, Formal analysis, Investigation

**Competing interests:** No competing interests declared ORCID 0000-0002-2245-8907

### Dawid Droździel

- Laboratory of Brain Imaging, Nencki Institute of Experimental Biology of the Polish Academy of Sciences, Warsaw, Poland

**Contribution:** Methodology, Project administration, Investigation

**Competing interests:** No competing interests declared

### Agnieszka Widlarz

- Chair of Rhythmics and Piano Improvisation, Department of Choir Conducting and Singing, Music Education and Rhythmics, The Chopin University of Music, Warsaw, Poland **Contribution**: Methodology

**Competing interests:** No competing interests declared

### Aleksandra M. Herman

- Laboratory of Brain Imaging, Nencki Institute of Experimental Biology of the Polish Academy of Sciences, Warsaw, Poland

**Contribution:** Conceptualization, Methodology, Investigation, Writing - review & editing, Supervision, Project administration.

**Competing interests:** No competing interests declared ORCID 0000-0002-3338-0543

### Artur Marchewka

- Laboratory of Brain Imaging, Nencki Institute of Experimental Biology of the Polish Academy of Sciences, Warsaw, Poland

**Contribution:** Conceptualization, Methodology, Resources, Writing - review & editing, Supervision, Project administration, Funding acquisition.

**Competing interests:** No competing interests declared ORCID 0000-0002-1982-3299

#### Ethics

This study was approved by the Research Ethics Committee at the Institute of Psychology of the Jagiellonian University, Kraków, Poland and the experiment has been carried out in accordance with The Code of Ethics of the World Medical Association (Declaration of Helsinki). All participants provided written informed consent and were compensated for their time.

## Data-availability statement

Group-level statistical maps and behavioural data, as well as the code used to perform visualisations can be found on the Open Science Platform: https://osf.io/q8abr

## Acknowledgements

This study was supported by the National Science Centre (Narodowe Centrum Nauki) grant number 2018/30/E/HS6/00206. AH is supported by the Foundation for Polish Science (FNP).

We would like to thank: all the participants for the effort and dedication they put into training; Ms Agnieszka Kulesza for her involvement in data acquisition; Mrs Maria Chełkowska-Zacharewicz for the advice on improvements in the experimental design, procedure and communication aimed at reducing participant’s stress during the final performance (TP4). Finally, we thank Mrs Katarzyna Kiwior for designing and running the piano training course, and her huge involvement in the learning process of the participants, which was essential for the study’s success.

[…] no matter how hard a thing is to do, once it has been done it’ll become a whole lot easier [..].

- Terry Pratchett. Maskerade

